# Neural differentiation is moderated by age in scene- but not face-selective cortical regions

**DOI:** 10.1101/2020.01.21.914432

**Authors:** Sabina Srokova, Paul F. Hill, Joshua D. Koen, Danielle R. King, Michael D. Rugg

## Abstract

The aging brain is characterized by neural dedifferentiation – an apparent decrease in the functional selectivity of category-selective cortical regions. Age-related reductions in neural differentiation have been proposed to play a causal role in cognitive aging. Recent findings suggest, however, that age-related dedifferentiation is not equally evident for all stimulus categories and, additionally, that the relationship between neural differentiation and cognitive performance is not moderated by age. In light of these findings, in the present experiment younger and older human adults (males and females) underwent fMRI as they studied words paired with images of scenes or faces prior to a subsequent memory task. Neural selectivity was measured in two scene-selective (parahippocampal place area and retrosplenial cortex) and two face-selective (fusiform and occipital face areas) regions of interest using both a univariate differentiation index and multivoxel pattern similarity analysis. Both methods provided highly convergent results which revealed evidence of age-related reductions in neural dedifferentiation in scene-selective but not face-selective cortical regions. Additionally, neural differentiation in the parahippocampal place area demonstrated a positive, age-invariant relationship with subsequent source memory performance (recall of the image category paired with each recognized test word). These findings extend prior findings suggesting that age-related neural dedifferentiation is not a ubiquitous phenomenon, and that the specificity of neural responses to scenes is predictive subsequent memory performance independently of age.

**Significance Statement:** Increasing age is associated with reduced neural specificity in cortical regions that are selectively responsive to a given perceptual stimulus category (age-related neural dedifferentiation), a phenomenon which has been proposed to contribute to cognitive aging. Recent findings reveal that age-related neural dedifferentiation is not present for all types of visual stimulus categories, and the factors which determine when the phenomenon arises remain unclear. Here, we demonstrate that scene- but not face-selective cortical regions exhibit age-related neural dedifferentiation during an attentionally-demanding task. Additionally, we report that higher neural selectivity in the scene-selective ‘parahippocampal place area’ is associated with better memory performance after controlling for variance associated with age group, adding to evidence that neural differentiation impacts cognition across the adult lifespan.

## 1. Introduction

Increasing age has been reported to be associated with reduced specificity and distinctiveness of neural representations, a phenomenon known as age-related neural dedifferentiation (for review, see Koen & Rugg, 2019; Koen et al., 2019, 2020). Computational models of cognitive aging suggest that neural dedifferentiation plays a role in age-related cognitive decline (Li et al., 2001; Li & Rieckmann, 2014). Specifically, the phenonemon has been proposed to arise from age-related reductions in neuromodulation, compromising the fidelity of neural representations (see also Abdulrahman et al, 2017).

In an early fMRI study of age-related neural dedifferentiation, Park et al. (2004), it was reported that healthy older adults demonstrated lower neural selectivity for preferred stimuli in voxels selective for four different perceptual categories (houses, chairs, pseudowords and faces). Although subsequent studies have reported convergent findings, the data suggest that age-related dedifferentiation is not ubiquitous. For example, whereas reduced neural selectivity is frequently reported in scene-selective (Carp et al. 2011; Voss et al., 2008; Zheng et al., 2018; Koen et al., 2019) and face-selective cortical regions (Park et al., 2004; Voss et al., 2008; Park et al., 2012), null findings for both of these classes of stimuli have also been reported (for scenes: Berron et al., 2018; for faces: Payer, et al., 2016). The evidence is also divergent for object and word stimuli. Although Park et al. (2004) reported age-related dedifferentiation for objects and orthographic stimuli, subsequent studies have found null age effects for both classes of stimuli (for objects: Chee et al., 2006; Zheng et al., 2018; Koen et al., 2019; for words: Voss et al., 2008). Additionally, Wang et al. (2016) reported null age effects on the accuracy of a multi-voxel classifier trained to discriminate neural activity elicited by words and pictures (see Abdulrahman et al., 2017, for similar null findings).

Metrics of neural differentiation have been reported to predict both memory performance for the experimental stimuli (e.g. Yassa et al., 2011; Berron et al., 2018; Bowman et al., 2019; Koen et al., 2019; Sommer et al., 2019; for related findings, see Du et al., 2016) as well as measures of cognitive performance derived from standardized tests (‘fluid’ processing in Park et al., 2010; ‘fluency’ in Koen et al., 2019). These findings are consistent with the possibility that age-related cognitive decline is driven by age differences in neural selectivity (dedifferentiation). Recent findings, coupled with a review of the relevant (though sparse) literature, suggest however that the relationship between neural differentiation and cognitive performance is age-invariant (Koen et al., 2019; Koen and Rugg, 2019). Although an age-invariant relationship between cognition and neural differentiation does not rule out a role for diminishing neural selectivity in mediating age-related cognitive decline, it does suggest that the contribution of neural selectivity to cognitive performance is stable across the lifespan.

In the present study, participants underwent fMRI while studying word-face and word-scene stimulus pairs. Memory was later assessed in two steps: recognition memory of the words, followed by recall of the associated image. Neural differentiation was operationalized both by a univariate differentiation index (Voss et al., 2008; Koen et al., 2019) and multi-voxel pattern similarity (Zheng et al., 2018; Koen et al., 2019; Sommer et al., 2019). Differentiation metrics were obtained from two face-selective (Fusiform face area, FFA; Occipital Face Area, OFA) and two scene-selective (Parahippocampal place area, PPA; Retrosplenial cortex, RSC) regions of interest (ROIs). Motivated by the recently reported finding of null age effects on neural selectivity for object stimuli (Koen et al., 2019), the current study aimed to examine whether the null effects of age extend to active processing of faces. Additionally, we aimed to replicate and extend the prior findings of age-related neural dedifferentiation for scene stimuli, and the relationships of neural differentiation with subsequent memory performance and psychometric measures of cognitive ability.

## 2. Materials and Methods

### 2.1 Ethics Statement

The experimental procedures described below were approved by The Institutional Review Boards of the University of Texas Southwestern Medical Center and the University of Texas at Dallas. All participants provided informed consent prior to taking part in the experiment.

### 2.2 Participants

Twenty-seven younger and 33 older adult volunteers were recruited from local communities surrounding The University of Texas at Dallas and the greater Dallas metropolitan area, and were compensated 30$/h. All volunteers were right-handed, had normal or corrected-to-normal vision, and were fluent English speakers before the age of five. Participants were excluded if they self-reported a history of cardiovascular or neurological disease, diabetes, substance abuse, use of medication affecting the central nervous system, or showed evidence of cognitive impairment based on their performance on a neuropsychological test battery (see below).

Three younger and three older adult participants were excluded from subsequent analyses for the following reasons: voluntary withdrawal from the study (N = 2), behavioral performance which resulted in not having enough trials (< 10) in a critical memory bin (N = 2), technical malfunction of the equipment (N = 1), and an incidental MRI finding (N = 1). Additionally, six older participants were excluded due to chance source memory performance, according to our pre-determined cutoff score (measure of source recollection, pSR < .1). The final sample therefore consisted of 24 young (age range: 18 – 28 years, 15 female) and 24 older adult (age range: 65 – 75 years, 14 female) participants. Demographic data and neuropsychological test performance are reported in Table 1.

**Table 1.** Demographic data and results of the neuropsychological test battery (mean, SD) for younger and older adults.

|  | **Younger Adults** | **Older Adults** | **p-value** |
| --- | --- | --- | --- |
| *N* | 24 | 24 |  |
| *Age* | 22.42 (3.24) | 70.00 (3.46) |  |
| *Years of Education* | 15.46 (2.65) | 16.71 (2.44) | NS |
| *MMSE* | 29.25 (0.90) | 29.33 (0.70) | NS |
| *CVLT Short Delay – Free* | 13.75 (2.00) | 11.88 (2.86) | 0.012 |
| *CVLT Short Delay – Cued* | 13.83 (2.32) | 13.08 (2.15) | NS |
| *CVLT Long Delay – Free* | 14.13 (2.11) | 12.79 (2.62) | NS |
| *CVLT Long Delay – Cued* | 14.38 (1.93) | 13.46 (2.13) | NS |
| *CVLT recognition – Hits* | 15.71 (0.46) | 15.25 (1.07) | NS |
| *CVLT recognition – False alarms* | 0.33 (0.70) | 1.67 (1.61) | 0.001 |
| *Logical Memory I* | 33.00 (4.76) | 28.00 (4.11) | < 0.001 |
| *Logical memory II* | 32.00 (4.80) | 25.83 (5.49) | < 0.001 |
| *SDMT* | 62.33 (11.27) | 49.29 (7.91) | < 0.001 |
| *Trails A (s)* | 20.20 (5.26) | 25.11 (6.46) | 0.006 |
| *Trails B (s)* | 44.12 (10.18) | 62.48 (16.77) | < 0.001 |
| *Digit Span Total* | 19.71 (4.14) | 18.79 (3.49) | NS |
| *Category fluency* | 23.71 (4.91) | 22.46 (5.35) | NS |
| *F-A-S* | 49.17 (12.85) | 46.29 (12.75) | NS |
| *WTAR* | 42.42 (3.46) | 44.54 (4.06) | NS |
| *Raven's* | 11.04 (0.86) | 9.50 (1.89) | 0.001 |
| *Speed Factor (RC1)* | -1.47 (2.01) | 1.56 (1.68) | < 0.001 |
| *Memory Factor (RC2)* | 1.53 (1.94) | -1.62 (2.42) | < 0.001 |
| *Crystallized intelligence Factor (RC3)* | 0.30 (1.42) | -0.39 (1.79) | NS |
| *Fluency Factor (RC4)* | 0.20 (1.18) | -0.10 (1.45) | NS |

*Digit Span total corresponds to the sum of forward and backward digit span.*
*Speed factor bears a negative number with better performance on tasks of processing speed.*
*NS = not significant*

Several of the participants in the present study had previously participated in one or more studies reported by our laboratory. Specifically, 4 older adults participated in the event related potential study reported by Koen et al. (2018), 1 older adult participated in a prior fMRI study reported by Koen et al. (2019), and 4 older adults took part in an fMRI experiment first reported by de Chastelaine et al. (2015).

### 2.3 Neuropsychological Testing

All participants completed our standard neuropsychological test battery consisting of the Mini-Mental State Examination (MMSE), The California Verbal Learning Test-II (CVLT; Delis et al., 2000), Wechsler Logical Memory Tests 1 and 2 (Wechsler, 2009), The Trail Making tests A and B (Reitan and Wolfson, 1985), the Symbol Digit Modalities test (SDMT; Smith, 1982), the F-A-S subtest of the Neurosensory Center Comprehensive Evaluation for Aphasia (Spreen and Benton, 1977), the Wechsler Adult Intelligence Scale–Revised subtests of forward and backward digit span (Wechsler, 1981), Category fluency test (Benton, 1968), Raven’s Progressive Matrices (List 1, Raven et al., 2000) and the Wechsler Test of Adult Reading (WTAR; Wechsler, 1981). Potential participants were excluded prior to the fMRI session if they scored < 27 on the MMSE, >1.5 SD below age norms on any standardized memory test, >1.5 SD below age norms on two or more standardized non-memory tests, or if their estimated full-scale IQ was < 100.

The neuropsychological test scores were reduced to four components based on the outcome of a principal components analysis applied to a prior large dataset from our laboratory. The dataset comprised scores from younger, middle aged and older adults (total N=154) (de Chastelaine et al. 2016). Four principal components with eigenvalues greater than 1, accounting for 64.1% of the variance, were retained and subjected to the Varimax rotation (Kaiser, 1958). The rotated components (RC) correspond roughly to processing speed (RC1), memory (RC2), crystallized intelligence (RC3), and fluency (RC4). The neuropsychological tests included in the analysis, their corresponding rotated factor weights, and the proportions of variance accounted for by the rotated components are presented in Table 2.

**Table 2:** Factor Loadings from the PCA, Varimax rotated, based on dataset previously reported by de Chastelaine et al. (2016).

|  | **Speed**<br>**(RC1)** | **Memory**<br>**(RC2)** | **Crystallized**<br>**Intelligence**<br>**(RC3)** | **Fluency**<br>**(RC4)** |
| --- | --- | --- | --- | --- |
| *CVLT composite* | -.19 | .84 | .08 | -.15 |
| *CVLT recognition – Hits* | -.20 | .42 | .23 | -.64 |
| *CVLT recognition – False alarms* | .21 | -.69 | .26 | -.17 |
| *Logical memory composite* | .10 | .67 | .18 | .02 |
| *Trails A (s)* | .91 | -.09 | -.05 | -.14 |
| *Trails B (s)* | .85 | -.09 | -.28 | .08 |
| *SDMT* | -.59 | .40 | .08 | .30 |
| *Digit Span* | -.16 | .01 | .80 | -.08 |
| *Category fluency* | -.34 | .23 | .14 | .63 |
| *F-A-S* | -.12 | .06 | .46 | .57 |
| *WTAR* | -.12 | .12 | .79 | .21 |
| *Raven’s* | -.33 | .48 | .10 | .05 |
| *Eigenvalue* | 3.65 | 1.70 | 1.28 | 1.06 |
| *Variance explained (before rotation)* | .20 | .14 | .11 | .09 |
| *Variance explained (after rotation)* | .19 | .19 | .15 | .11 |

### 2.4. Experimental Materials and Procedure

#### 2.4.1. Experimental Procedure and Materials

Experimental stimuli were presented using Cogent 2000 software (www.vislab.ucl.ac.uk/cogent_2000.php) implemented in Matlab 2012b (www.mathworks.com). The stimuli were projected onto a translucent screen attached at the rear of the MRI bore and were viewed through a mirror mounted on the scanner head coil. Participants completed two study-test cycles inside the scanner. For each cycle, study and test phases were each split into two scanning sessions, with a 30s rest period midway through each session. The critical experimental stimuli were distributed across four study and four test sub-lists, with a single sub-list per scanning session. Therefore, participants’ memory for the first two study sub-lists was tested in two memory test sessions before continuing to the second cycle. The critical stimuli comprised 288 concrete nouns, 96 colored images of male and female faces (face stimuli obtained from Minear & Park (2004) database), and 96 colored images of urban and rural scenes.

All images of faces and scenes were scaled at 256 x 256 pixels. An additional 68 words and 40 images were used as fillers at the beginning of each scan session and immediately after each break or as practice stimuli. The critical stimuli were interspersed with 96 null trials (white fixation cross) in both the study and test lists (24 trials per sub-list). Stimuli were selected randomly without replacement to create twenty-four different stimulus sets for yoked younger and older adult pairs. Study and test trials were pseudorandomized such that participants were not presented with more than three consecutive trials belonging to the same image class, or more than two consecutive null trials.

#### 2.4.2. Study Phase

Participants completed two scanned study-test cycles. Each cycle included two study blocks. The blocks each contained 24 null trials and 48 critical words, half of which were paired with an image of a face and a half paired with a scene image. The word was presented in the upper half of the screen with the image beneath it and a white fixation cross positioned between the two items (see Figure 1). Words were presented in a white font 30pt uppercase Helvetica over a black background. A study trial began with a red fixation cross for a duration of 500ms, followed by the presentation of the word-image pair for 2000ms. This was followed a white fixation cross for a further 2000ms. When a word was paired with a face, the instructions were to imagine the person depicted by the image interacting with the object denoted by the word. For word-scene trials, the task was to imagine a scenario in which the object denoted by the word is interacting with elements of the scene. To ensure adherence to task instructions, participants were asked to rate the vividness of each imagined scenario on a three-point scale: ‘Not vivid, ‘Somewhat vivid’, to ‘Very vivid’. Responses were recorded with right-hand index, middle and ring fingers using a scanner-compatible button box. Only trials on which ratings were made between 450-4500ms post-stimulus onset were included in the analyses described below. Trials attracting multiple responses were excluded from behavioral analyses and included as events of no interest in the fMRI analyses.

**Figure 1:**
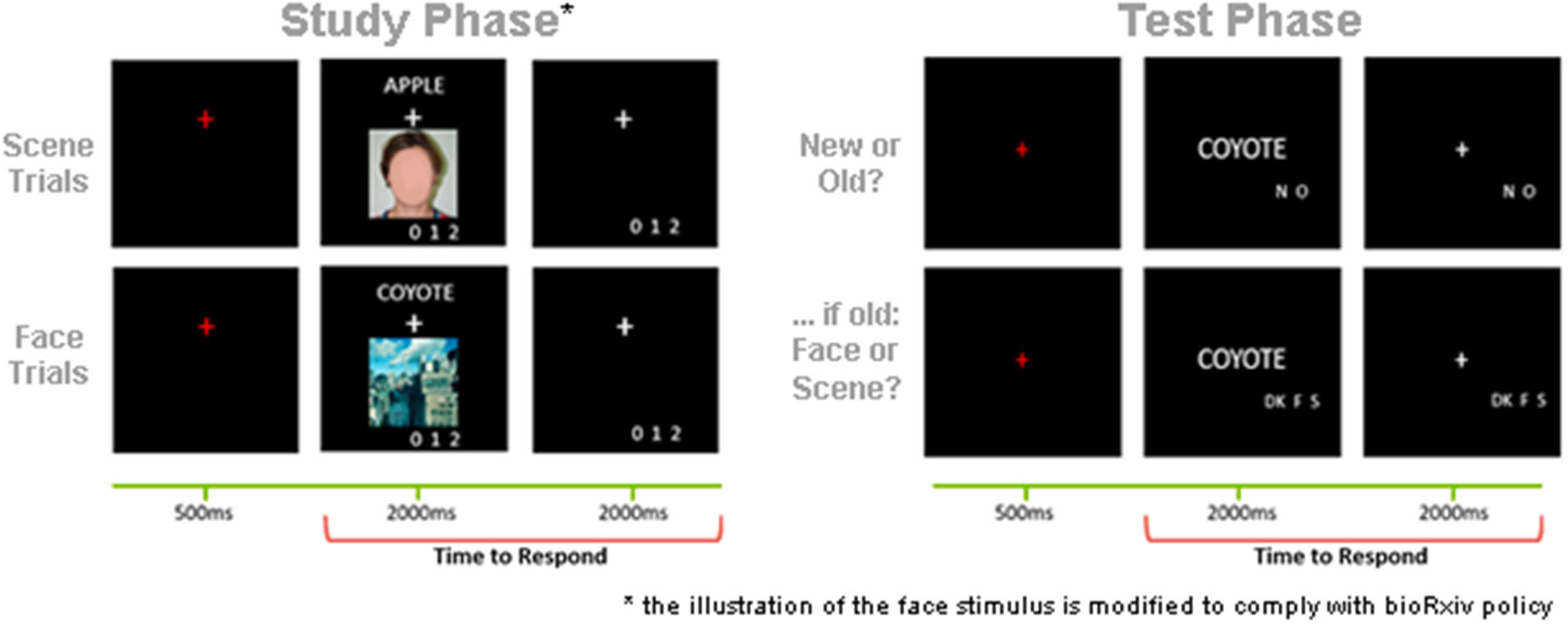
Overview of the encoding task and subsequent memory test. At encoding, participants were asked to “Imagine the person interacting with the object denoted by the word.” (face trials) or to “Imagine the object denoted by the word interacting with the scene.” (scene trials).

#### 2.4.3. Test Phase

The test phase was also conducted within the fMRI scanner (the fMRI data will be reported in a separate communication). While undergoing scanning, participants’ memory for the studied items was tested across two test lists (two sub-lists per study-test cycle). Each sub-list comprised 48 studied words, 24 new words, and 24 null trials. Each test trial began with a 500ms duration red fixation cross, followed by the test word, which was presented for 2000ms, and a white fixation cross for 2000ms. Participants were required to indicate whether they remembered studying the test words by making an ‘Old’ or a ‘New’ judgment. Instructions were to respond ‘Old’ only if they were confident the word had been studied. For test items endorsed ‘Old’, participants were prompted to make a source memory judgement, during which they signaled whether the word had been studied along with a face or a scene. An additional ‘Don’t Know’ response option was available to discourage guessing. The source memory prompt was presented immediately after the ‘Old’/’New’ memory response had been made. Test items receiving a ‘New’ judgement were followed by a 2000ms duration white fixation cross. Test responses were made with the index, middle and ring fingers of the right hand on a scanner-compatible button box. The buttons were counterbalanced across participants such that the ‘Old’/’New’ judgment were made with the index and middle finger, while the source judgements were counterbalanced across the index, middle, and ring fingers with the constraint that the ‘Don’t know’ response was never assigned to the middle finger. Similarly to the study phase, trials associated with responses made outside of a 500ms– 4500ms post-stimulus window were not considered in the analyses and were included as events of no interest.

### 2.5. Data Acquisition and Analysis

#### 2.5.1 Experimental Design and Statistical Analysis

The main independent variables in the analyses described below include age group (younger vs older adults), image category of the study trials (faces vs scenes), and the two face-selective and two scene-selective regions-of-interest (ROIs): Fusiform Face Area (FFA) and Occipital Face Area (OFA) as face-selective ROIs; Parahippocampal Place Area (PPA) and Retrosplenial cortex (RSC) as scene-selective ROIs.

Statistical analyses were conducted using R Software (R Core Team, 2019) and all tests were considered significant at p < 0.05. Analyses of variance were performed using the *afex* package (Singmann et al., 2016) and the degrees of freedom were corrected for nonsphericity using the Greenhouse and Geisser (1959) procedure. All *t*-tests were performed as Welch’s unequal variance tests using the t-test function in base R. Effect sizes are reported as partial-*η*^2^ for the analysis of variance (ANOVA) results and the package *effsize* (Torchiano, 2019) was used for Cohen’s d in pairwise comparisons (Cohen, 1988). Linear regression models were employed using the lm function in base R, and partial correlations were conducted using the function pcor.test in the *ppcor* package (Kim, 2015). Principal components analysis (Hotelling, 1933; Abdi and Williams, 2008) on the neuropsychological test scores was implemented with the *psych* package (Revelle, 2017).

#### 2.5.2. Behavioral Data Analysis

Study and test trials were binned according to their subsequent memory status. We focused on item recognition performance as reflected in the initial ‘Old’ / ‘New’ judgement, and source memory performance as indexed by the subsequent ‘Scene’/ ‘Face’ / ‘Don’t Know’ judgement. Trials that received no response or multiple responses were excluded. Item Memory performance was computed as the difference between the overall hit rate and the false alarm rate:

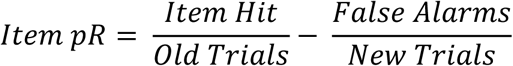

The hit rate was calculated as the proportion of trials which were correctly endorsed as ‘Old’ relative to the total number of old trials, regardless of their subsequent source memory judgement. The false alarm rate was calculated as the proportion of new trials incorrectly endorsed as ‘Old’ relative to all new trials. The overall item recognition accuracy was submitted to a 2 (Age group) x 2 (Image class) mixed factorial ANOVA.

Additionally, source memory accuracy was computed using a modified single high-threshold model (Snodgrass and Corwin, 1988) according to the following formula (see Gottlieb et al., 2010; Mattson et al. 2014):

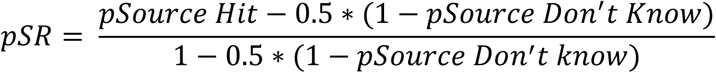

where ‘pSource Hit’ refers to the proportion of correctly recognized test items endorsed with a correct source memory judgement at test and ‘pSource Don’t Know’ refers to items that were correctly recognized but received a ‘Don’t Know’ source memory response. Given the design of this experiment, our source memory metric necessarily encompasses both face and scene trials. Therefore, we collapsed source memory performance across image type and compared performance between the two age groups with an independent samples t-test.

Other behavioral measures included reaction time (RT) and vividness ratings for the encoding trials. RT was calculated as the median time to make a vividness rating. Both RTs and the vividness ratings were computed separately for trials corresponding to each image class and binned according to whether or not they were associated with a correct source judgment at test. The vividness ratings and RTs were submitted to separate 2 (Age group) x 2 (Image class) x 2 (Subsequent memory) mixed factorial ANOVAs.

#### 2.5.3. MRI Data Acquisition and Preprocessing

Functional and structural MRI data were acquired at 3T using a Philips Achieva MRI scanner (Philips Medical Systems, Andover, MA) equipped with a 32 channel receiver head coil. The functional scans were acquired with a T2*-weighted, blood-oxygen level-dependent echoplanar imaging (EPI) sequence (sensitivity encoding [SENSE] factor = 2, flip angle = 70°, 80 x 78 matrix, field of view [FOV] = 24 cm, repetition time [TR] = 2000 ms, and echo time [TE] = 30 ms). EPI volumes comprised 34 slices (1mm interslice gap) at a voxel size of 3×3×3 mm, acquired in an ascending order and parallel to the anterior-posterior commissure line. Structural images were obtained with a T1-weighted MPRAGE sequence (FOV = 240 x 240, 1×1×1 mm isotropic voxels, sagittal acquisition).

MRI data were preprocessed and analyzed using a combination of Statistical Parametric Mapping (SPM12, Wellcome Department of Cognitive Neurology, London, UK) and custom Matlab scripts. The functional images were realigned to the mean EPI image and slice-time corrected using sinc interpolation to the middle slice. The images were then subjected to reorientation and spatial normalization with respect to a sample-specific template following previously published procedures (de Chastelaine et al. 2011, 2016). Functional images were smoothed with an 8 mm full-width half maximum Gaussian kernel prior to region-of-interest (ROI) selection. Estimation of differentiation indices and PSA were conducted on unsmoothed data.

#### 2.5.4. MRI Data Analysis

The analyses reported here focus on the data from the study sessions (analyses of the test data will be reported in a separate paper). The ROIs were derived from univariate fMRI analyses across the four study sessions, which were performed in two stages. In the first stage, separate GLMs were constructed for each participant by sorting the study trials into two categories depending on the trial type: scene trials and face trials. Trials belonging to each of these categories were modeled with a 2s duration boxcar function onsetting concurrently with the onset of the study word-image pair, convolved with a canonical hemodynamic response function (HRF). Filler trials, null trials, and trials which received multiple or no responses were modeled as covariates of no interest. Additional covariates of no interest included the 30s duration rest periods midway through each study session and the six regressors representing motion-related variance (three representing rigid-body translation and three for rigid-body rotation along the three axes). Trials with translational displacement greater than 1mm or with rotational displacement greater than 1° in any direction were modeled as covariates of no interest and hence removed from the analysis. In the second stage, the parameter estimates of the two events of interest were carried over to a second-level random effects 2 x 2 factorial ANOVA with age (younger, older) treated as the between-subjects factor, and trial type (scene, face) as the within-subjects factor.

For the purposes of the differentiation index analyses and the PSA, the unsmoothed data from each of the four total study sessions were concatenated using the *spm_fmri_concatenate* function and subjected to a ‘least-squares-all’ analysis (Rissman et al., 2004; Mumford et al., 2014) to estimate the BOLD response for each trial. Each event was modeled with a 2s duration boxcar function and convolved with a canonical HRF. The covariates of no interest included the 6 motion regressors described above and the four session specific means.

#### 2.5.5. Region-of-Interest Selection

Two face-selective (FFA, OFA) and two scene-selective (PPA, RSC) ROIs were empirically defined via a second-level GLM that contrasted scenes and faces, (thresholded at p < .01 (uncorrected)) across all participants without regard to the factor of age group. The contrasts were inclusively masked with the ‘Neuromorphometrics’ atlas provided in SPM12. The face > scene contrast was masked with the atlas’s fusiform gyrus and parahippocampal gyrus to derive the FFA mask, and the OFA was defined by inclusively masking the contrasts with inferior occipital and occipital fusiform gyri. The scene > face contrast was masked with the fusiform and parahippocampal gyri to identify the PPA. As Neuromorphometrics does not provide a mask for the RSC, we searched the Neurosynth database using the term “retrosplenial” (search in August 2019, search results FDR-corrected at p < .00001; Yarkoni et al., 2011) and used the outcome to create the RSC mask. All ROIs were collapsed across the two hemispheres.

#### 2.5.6. Differentiation Index

We computed a differentiation index for each ROI as a measure of the selectivity of neural responses at the regional level (Voss et al., 2008; Koen et al., 2019). The differentiation index for a given ROI was computed as the difference between the mean BOLD response for trials of a preferred stimulus class and the mean BOLD response for trials of the non-preferred class, divided by pooled standard deviation:

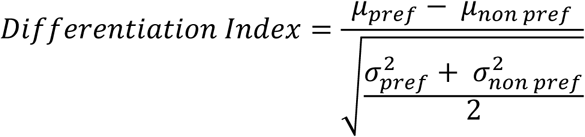

Therefore, a higher differentiation index indicates greater selectivity for a given ROI (note that because of the scaling function, the differentiation index is insensitive to individual or group differences in the gain of the hemodynamic response function mediating between neural activity and the fMRI BOLD response). We computed a differentiation index for each of the four ROIs for each participant. The resulting indices were subjected to a 2 (Age group) x 4 (ROI) mixed factorial ANOVA (see results). We conducted an additional ANOVA of the differentiation indices computed only from the trials that went on to receive a source correct memory response. The goal of this additional analysis was to ascertain whether any age differences arising from the original analysis were a reflection of the differential mixing of trial types as a function of age group (on average, young participants had a higher proportion of source correct study trials than did older adults).

Neural dedifferentiation may manifest as a reduced neuronal response to a preferred stimulus category (i.e. neural attenuation), as an elevated response to a non-preferred category (i.e. neural broadening), or as the combination of both phenomena (Park et al., 2012; Koen & Rugg, 2019). The differentiation index is insensitive to this distinction. Thus, we also examined the *β*-parameters, averaged across all voxels within each ROI, reflecting responses to scene and face trials in ROIs where significant age differences in neural differentiation were identified. The *β*-parameters were subjected to a 2 (Age group) x 2 (ROI) x 2 (Image class) mixed-factorial ANOVA.

Finally, to examine whether neural differentiation predicted memory performance, for each ROI we constructed regression models that employed differentiation index and age-group as predictor variables, and, in parallel models, either source or item memory performance as the dependent variable. Initial versions of the models also included the interaction between differentiation index and age group as an additional predictor variable. In no case did the interaction term account for a significant fraction of the variance in performance (p > .05). Results are reported below for the reduced models that excluded the interaction term.

#### 2.5.7. Multivoxel Pattern Similarity Analysis

Multivoxel pattern similarity analysis (PSA) was conducted in a similar fashion to Koen et al. (2019) to complement the univariate analyses described above. The similarity measures were derived from single-trial, voxel-wise *β*-parameters (see Methods 2.5.4 above). For each participant and ROI, we first computed a within-category similarity metric. This was achieved by computing the correlations across voxels between each study trial and all other study trials belonging to the same image category, subjecting the resulting correlations to a Fisher’s z transformation, and averaging them. The between-category similarity was calculated in an analogous fashion except that the correlations were estimated across rather than within image category. The between- and within-similarity was always computed across trials of different scanning sessions to avoid potential bias arising from carry-over effects (Mumford et al. 2014). The similarity index was then computed as the difference between the within- and between-category similarity metrics. This index can be used as a metric of neural differentiation as it reflects the extent to which different perceptual categories evoke consistent patterns of neural responses within a given region of interest. As in the case of the differentiation index described above, this correlation-based metric is insensitive to individual differences in hemodynamic gain.

The similarity indices were subjected to a 2 (Age group) x 4 (ROI) mixed factorial-ANOVA. As with the analyses of the differentiation indices, we also computed pattern similarity separately for trials that went on to receive a source correct memory response. Additionally, similarity indices were employed in regression analyses aimed at predicting behavioral performance. These analyses were exactly analogous to those conducted on the differentiation indices.

## 3. Results

Demographic data and the outcomes of the neuropsychological test battery are presented in Table 1. The groups were well-matched for years of education and MMSE but showed a typical pattern of age-related differences in cognitive performance. Thus, relative to the older group, younger adults had better performance on a subset of declarative memory tests, including the CVLT short free recall test and the logical memory subtests of the WMS. The younger adults also made significantly fewer recognition false alarms on the CVLT recognition memory test, and out-performed the older group on the speeded tests (Trails A, Trails B, and Symbol Digit Modalities) and Raven’s progressive matrices.

The rotated factor loadings (see Methods) were applied to each participant’s neuropsychological test scores, and the resulting factor scores for the four rotated components are presented at the bottom of Table 2. Consistent with the individual neuropsychological tests, there were age differences in the Speed and the Memory constructs. There were no age differences in the Crystallized Intelligence or Fluency factors.

### 3.2. Behavioral Results

#### 3.2.1. Study Performance

Mean study reaction times (RTs) and vividness ratings are reported in Table 4, separated by image category and age group. A 2 (Age group) x 2 (Image category) x 2 (Memory: source correct vs. source incorrect/don’t know and item misses) mixed factorial ANOVA on the RT data revealed a significant main effect of category, reflecting faster responses in face trials (F_(1,46)_ = 5.350, p = .025, partial-*η*^2^ = 0.101), but the remaining main effects and all interactions were not significant (ps > 0.1). A 2 (Age group) x 2 (Image category) x 2 (Memory) ANOVA on the mean vividness ratings revealed a significant main effect of memory (trials rated as more vivid were associated with better source memory performance), (F_(1,46)_ = 53.436, p < 0.001, partial-*η*^2^ = 0.537). There was no effect of age (F_(1,46)_ = 3.120, p = 0.084, partial-*η*^2^ = 0.064), category (F_(1,46)_ = 0.656, p = 0.409, partial-*η*^2^ = 0.015), and no interaction effects (ps > 0.18).

**Table 3:** The voxel size and peak MNI coordinates for each ROI

|  | Voxel Size | Peak MNI Coordinates |  |  |
| --- | --- | --- | --- | --- |
|  |  | X | Y | Z |
| *R. Occipital Face Area* | 98 | 45 | -79 | -16 |
| *L. Occipital Face Area* | 24 | -45 | -85 | -10 |
| *R. Fusiform Face Area* | 34 | 45 | -43 | -28 |
| *L. Fusiform Face Area* | 10 | -42 | -49 | -25 |
| *R. Parahippocampal Place Area* | 219 | 30 | -40 | -19 |
| *L. Parahippocampal Place Area* | 249 | -27 | -46 | -16 |
| *R. Retrosplenial Cortex* | 168 | 18 | -58 | 14 |
| *L. Retrosplenial Cortex* | 211 | -15 | -61 | 11 |

**Table 4.** Mean (SD) Study phase performance in younger and older adult groups.

|  | Young Adults |  | Older Adults |  |
| --- | --- | --- | --- | --- |
|  | Faces | Scenes | Faces | Scenes |
| **Vividness Ratings** |  |  |  |  |
| Source Correct Memory | 2.42 (.32) | 2.44 (.32) | 2.24 (.39) | 2.18 (.43) |
| Incorrect Memory | 2.23 (.42) | 2.13 (.51) | 2.06 (.46) | 2.01 (.49) |
| **Reaction Time (ms)** |  |  |  |  |
| Source Correct Memory | 2369 (678) | 2398 (628) | 2130 (570) | 2266 (524) |
| Incorrect Memory | 2351 (658) | 2350 (633) | 2285 (605) | 2327 (579) |

#### 3.2.2. Memory Performance

Memory performance on the experimental task is summarized in Table 5. A 2 (Age group) x 2 (Image category) mixed factorial ANOVA on item recognition identified a significant main effect of image category (F_(1,46)_ = 5.443, p = 0.024, partial-*η*^2^ = 0.106), and a main effect of age group (F_(1,46)_ = 10.112, p = 0.003, partial-*η*^2^ = 0.180). There was no significant interaction between the two factors (F_(1,46)_ = 0.766, p = 0.386, partial-*η*^2^ = 0.016). The main effect of image class reflected higher item memory performance for words paired with faces relative to scenes. Additionally, overall item recognition performance was significantly greater for younger than older adults. An independent samples t-test on source memory performance (pSR) revealed a significant difference in favor of the younger group (t_(45.12)_ = 3.440, p = 0.001, d = 1.01).

**Table 5.**
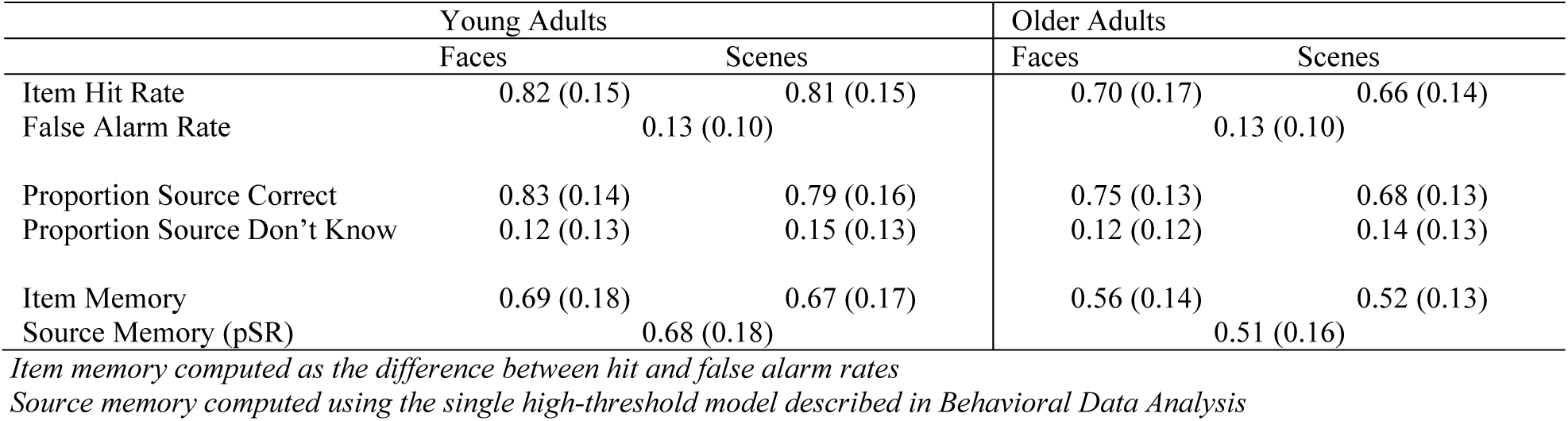
Mean (SD) Item and Source memory performance for younger and older adult groups.

#### 3.3.1. fMRI Differentiation Index

The differentiation indices were subjected to a 2 (age group) x 4 (ROI) mixed factorial ANOVA. The ANOVA revealed a main effect of ROI (F_(2.11, 96.87)_ = 29.498, p < 0.001, partial-*η*^2^ = 0.391), a main effect of age group (F_(1, 46)_ = 7.389, p = 0.009, partial-*η*^2^ = 0.138), and a significant age-by-ROI interaction (F_(2.11, 96.87)_ = 9.025 p < 0.001, partial-*η*^2^ = 0.164). Two follow-up ANOVAs were performed separately for the face-selective and scene-selective ROIs. The 2 (age group) x 2 (scene-selective ROIs) ANOVA resulted in a significant main effect of ROI (F_(1, 46)_ = 115.71, p < 0.001, partial-*η*^2^ = 0.715), a significant main effect of age group (F_(1, 46)_ = 24.006, p < 0.001, partial-*η*^2^ = 0.343), and a near-significant age-by-ROI interaction (F_(1, 46)_ = 3.869, p = 0.055, partial-*η*^2^ = 0.078). As is illustrated in Figure 3.A, the main effect of age group is driven by reduced neural differentiation in the older age group in both ROIs: PPA (t_(45.50)_ = 4.693, p < 0.001, d = 1.355), and RSC (t_(45.95)_ = 3.763, p < 0.001, d = 1.086). An analogous 2 (age group) x 2 (face-selective ROIs) ANOVA resulted in only a weak trend toward an age-by-ROI interaction (F_(1, 46)_ = 3.679, p = 0.061, partial-*η*^2^ = 0.074), and no main effect for ROI (F_(1, 46)_ = 0.637, p = 0.429, partial-*η*^2^ = 0.014), or age group (F_(1, 46)_ = 0.265, p = 0.609, partial-*η*^2^ = 0.006). Unsurprisingly, therefore, there were null effects of age on neural differentiation in both FFA (t_(45.81)_ = 0.401, p = 0.690), and OFA (t_(42.92)_ = −1.381, p = 0.175). Each of the differentiation indices illustrated in Figure 3.A differed significantly from zero in both age groups (ps < 0.002). Together, these results indicate that age group moderated neural differentiation within the scene-selective but not the face-selective ROIs.

**Figure 2:**
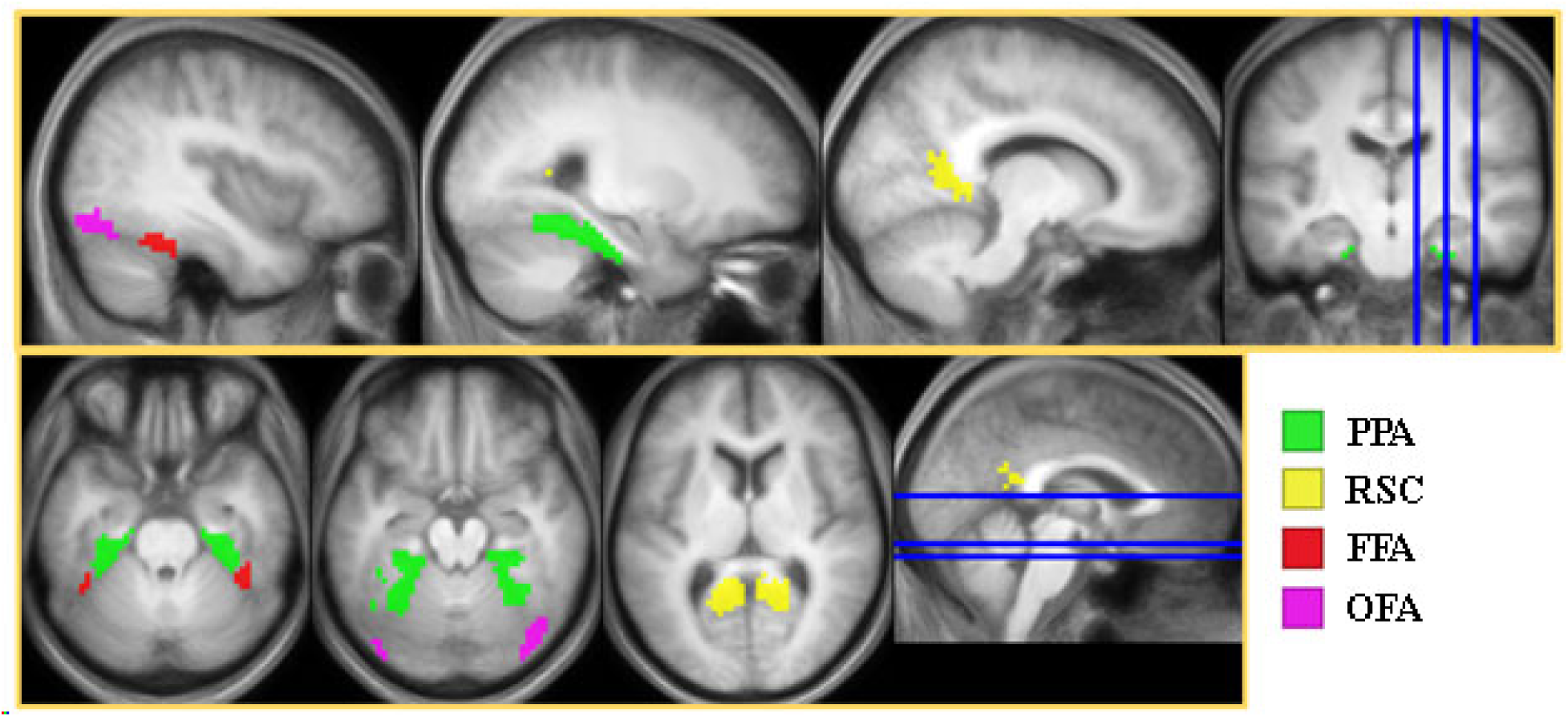
Bilateral scene- and face-selective ROIs derived using a second-level GLM contrasting faces and scenes, inclusively masked with Neuromorphometrics in SPM (PPA, FFA, OFA) or with Neurosynth (RSC).

**Figure 3:**
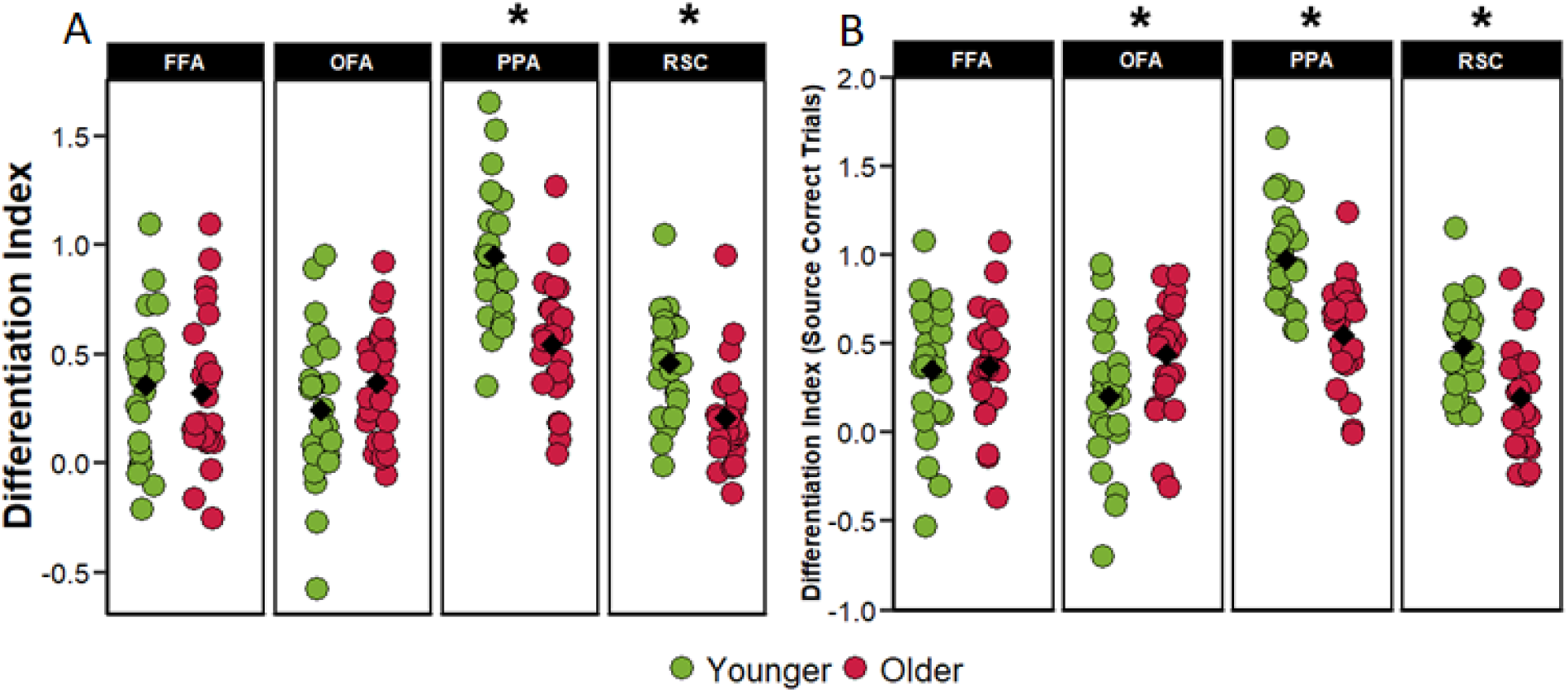
(A) Univariate differentiation indices computed collapsing across all trials regardless of subsequent memory performance. (B) Differentiation indices computed for only those trials that went on to receive a source-correct response at subsequent retrieval.

In a follow-up analysis, the differentiation index was computed separately for stimulus pairs according to whether they went on to receive a correct or any from incorrect response during a subsequent memory task. A 2 (age group) x 4 (ROI) x 2 (memory Status) mixed factorial ANOVA revealed a main effect of ROI (F_(2.09, 96.21)_ = 23.511, p < . 001, partial-*η*^2^ = .338), a main effect of age group (F_(1, 46)_ = 6.737, p = .013, partial-*η*^2^ = . 128), a significant age-by-ROI interaction (F_(2,09)_ = 6.250, p = .002, partial-*η*^2^ = .119), and a three-way interaction between age, ROI and memory status (F_(1.81, 83.16)_ = 4.483, p = .017, partial-*η*^2^ = .089). However, the analysis did not identify a main effect of memory (F_(1, 46)_ = 1.714, p = .197, partial-*η*^2^ = .036), nor a memory-by-age or memory-by-ROI interaction (F_(1, 46)_ = 2.567, p = .116, partial-*η*^2^ = .052, and F_(1.81, 83.16)_ = .605, p = . 532, partial-*η*^2^ = .013, respectively). Pairwise follow-up tests failed to identify significant differences between differentiation indices computed separately for the two classes of subsequent memory judgment in any of the ROIs in either age group (ps > 0.178).

We went on the examine the differentiation indices only for trials that were later associated with a source-correct memory response to ensure that the age-differences reported above were not driven by the differential mixing of source correct and source incorrect trials (given the age differences in source memory, see Methods 2.5.6). The ANOVA identified a significant main effect of ROI (F_(1.89, 86.74)_ = 22.40, p < 0.001, partial-*η*^2^ = 0.327), a main effect of age group (F_(1, 46)_ = 4.89, p = 0.032, partial-*η*^2^ = 0.096), and an Age-by-ROI interaction (F_(1.89, 86.74)_ = 11.103, p < 0.001, partial-*η*^2^ = 0.194). Similarly to the analyses of study trials collapsed across memory performance, we followed up the significant ROI by age group interaction with subsidiary 2 (age group) x 2 (face-selective ROIs) and 2 (age group) x 2 (scene-selective ROIs) ANOVAs. In the scene-selective regions, we identified a significant main effect of age-group (F_(1, 46)_ = 22.921, p < .001, partial-*η*^2^ = .333), a main effect of ROI (F_(1, 46)_ = 133.684, p < 0.001, partial-*η*^2^ = 0.744), but only a trend towards an age-by-ROI interaction (F_(1, 46)_ =3.938, p = 0.053, partial-*η*^2^ = 0.079). As evident in Figure 3.B, the effects of age on neural differentiation within the scene-selective regions were characterized by reduced differentiation indices in both PPA (t_(45.98)_ = 5.281, p < 0.001, d = 1.524), and RSC (t_(44.79)_ = 3.359, p = 0.002, d = 0.970). The analogous analysis in the face-selective regions revealed a significant age-by-ROI interaction (F_(1, 46)_ = 4.172, p = 0.047, partial-*η*^2^ = 0.083), but the ANOVA did not reveal main effects of age or ROI (F_(1, 46)_ = 2.013, p = 0.163, partial-*η*^2^ = 0.042 and F_(1, 46)_ = 0.640, p = 0.428, partial-*η*^2^ = 0.014, respectively). Subsequent pairwise comparisons demonstrated significantly greater differentiation in older relative to younger adults in the OFA (t_(43.92)_ = −2.204, p = 0.032, d = 0.636), but no age differences in the FFA (t_(44.94)_ = −0.258, p = 0.797, d = 0.075). As in the prior analyses, each of the differentiation indices illustrated in Figure 3.B was significantly different from zero in both age groups (ps < 0.019). Overall, restricting analyses to only those encoding trials receiving a subsequent source correct response led to convergent results in scene-selective ROIs, whereby older adults demonstrated lower neural selectivity relative to younger adults.

To further examine age-related dedifferentiation effects in scene-selective regions, we examined whether reduced neural selectivity in older adults resulted from a reduction in BOLD signal for the preferred image category (neural attenuation) or an increase in BOLD signal to the non-preferred category (neural broadening). A 2 (Age group) x 2 (ROI) x 2 (Image class) mixed factorial ANOVA on the extracted β-parameters revealed a significant main effect of ROI (F_(1, 46)_ = 125.677, p < 0.001, partial-*η*^2^ = 0.732), and a main effect of stimulus category (F_(1, 46)_ = 223.252, p < 0.001, partial-*η*^2^ = 0.829), but a null effect of age group (F_(1, 46)_ = 0.591, p = 0.445, partial-*η*^2^ = 0.013), and a null age group-by-ROI interaction (F_(1, 46)_ = 0.032, p = 0.859, partial-*η*^2^ = 0.001). However, the ANOVA revealed a 2-way interaction between stimulus category and age group (F_(1, 46)_ = 25.859, p < 0.001, partial-*η*^2^ = 0.360), and ROI (F_(1, 46)_ = 65.59, p < 0.001, partial-*η*^2^ = 0.588). The 3-way interaction was not significant (F_(1, 46)_ = 1.553, p = 0.219, partial-*η*^2^ = 0.033). As is evident from Figure 4, there was an attenuated BOLD response to scenes in older participants across both scene ROIs (t_(44.94)_ = −2.894, p = 0.005, d = −0.591), accompanied by an elevated response to face stimuli (t_(44.94)_ = 2.659, p = 0.009, d = 0.543). Thus, age-related neural dedifferentiation in the scene-selective ROIs was driven by a combination of attenuated BOLD response to scenes and increased responses to faces.

**Figure 4.**
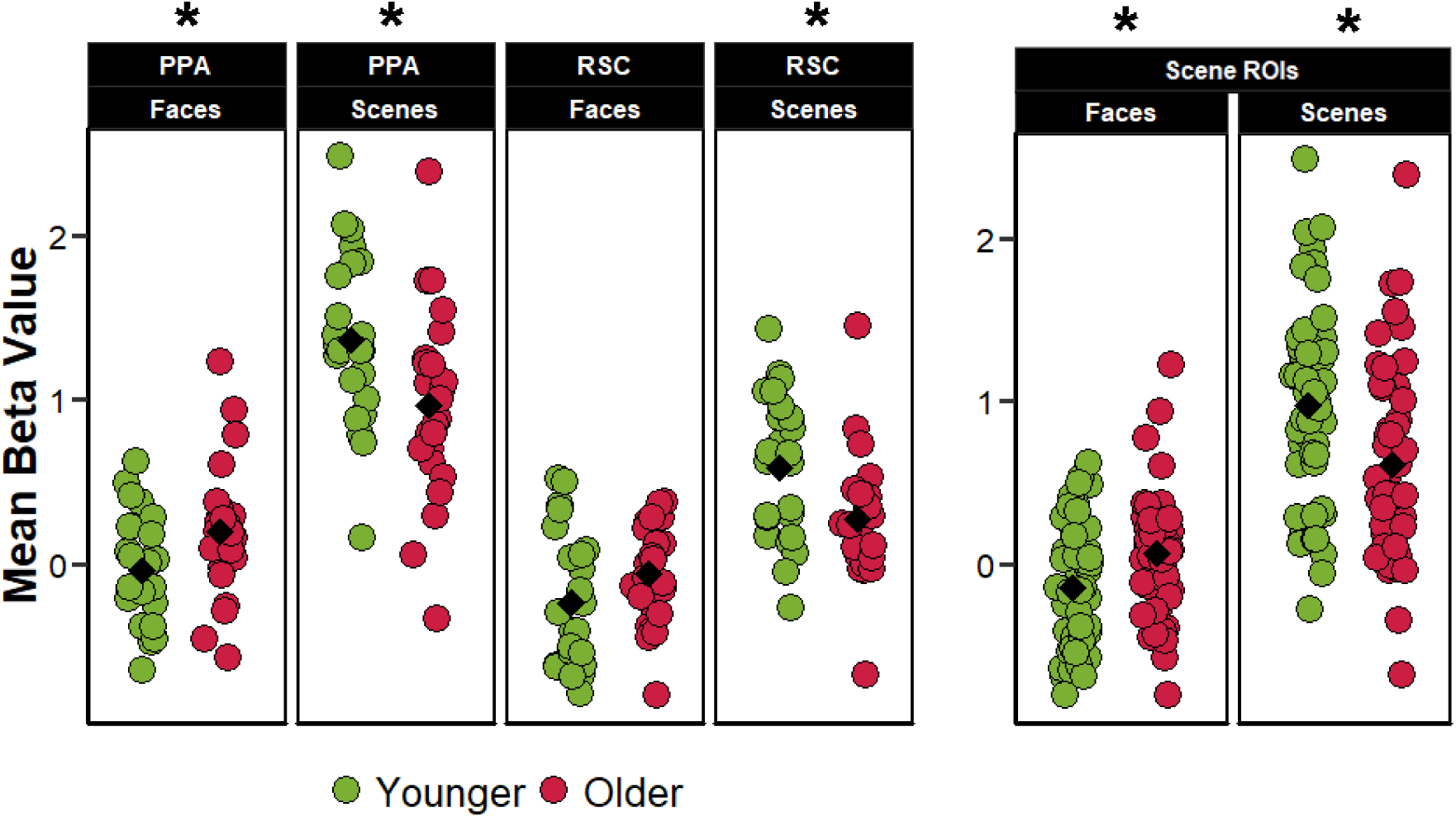
Across-trial mean β-parameters for face and scene trials in the scene-selective ROIs.

#### 3.3.2. Pattern Similarity Analysis

Multivoxel PSA (Kriegeskorte et al., 2008) was employed as a complement to the analysis of the differentiation index described above. We computed a within – between similarity metric in each ROI as an index of selectivity to the ROI’s preferred relative to the non-preferred stimulus class (see Methods). Analogous to the analyses of the differentiation index, the initial 2 (age group) x 4 (ROI) mixed factorial ANOVA was employed on the within – between similarity indices computed across all trials regardless of subsequent memory status. This revealed significant main effects of ROI (F_2.35, 108.24)_ = 11.924, p < 0.001, partial-*η*^2^ = 0.206), and age group (F_(1, 46)_ = 12.855, p < 0.001, partial-*η*^2^ = 0.218), along with a significant two-way interaction (F_(2.35, 108.24)_ = 4.981, p = 0.006, partial-*η*^2^ = 0.098). A subsequent 2 (age group) x 2 (ROI) mixed ANOVA focusing on just the scene-selective ROIs yielded a significant main effect of ROI (F_(1, 46)_ = 71.02, p < 0.001, partial-*η*^2^ = 0.607), a main effect of age (F_(1, 46)_ = 20.273, p < 0.001, partial-*η*^2^ = 0.306), and a significant age-by-ROI interaction (F_(1, 46)_ = 19.077, p < 0.001, partial-*η*^2^ = 0.293). An analogous 2 (age group) x 2 (ROI) ANOVA on the data from the face-selective ROIs failed to identify a significant age x ROI interaction (F_(1, 46)_ = 0.191, p = 0.174, partial-*η*^2^ = 0.040), nor did it reveal significant main effects of ROI (F_(1, 46)_ = 0.575, p = 0.452, partial-*η*^2^ = 0.012), or age group (F_(1, 46)_ = 3.091, p = 0.085, partial-*η*^2^ = 0.063). Follow-up pairwise comparisons examining age differences in each of the four ROIs revealed significantly lower similarity metrics for scenes in both the PPA (t_(40.50)_ =5.191, p < 0.001, d = 1.498), and RSC (t_(37.66)_ = 2.290, p = 0.027, d = 0.660). We did not however detect any age differences in similarity indices for faces within face-selective ROIs: FFA (t_(33.06)_ = 1.939, p = 0.061, d = 0.560), OFA (t_(45.46)_ = 0.626, p = 0.534, d = 0.181) (Figure 5.A). The similarity indices differed significantly from zero in all ROIs in both age groups (ps < 0.001). These results indicate that, when computed across all encoding trials, within – between pattern similarity was moderated by age in the scene- but not the face-selective ROIs.

**Figure 5:**
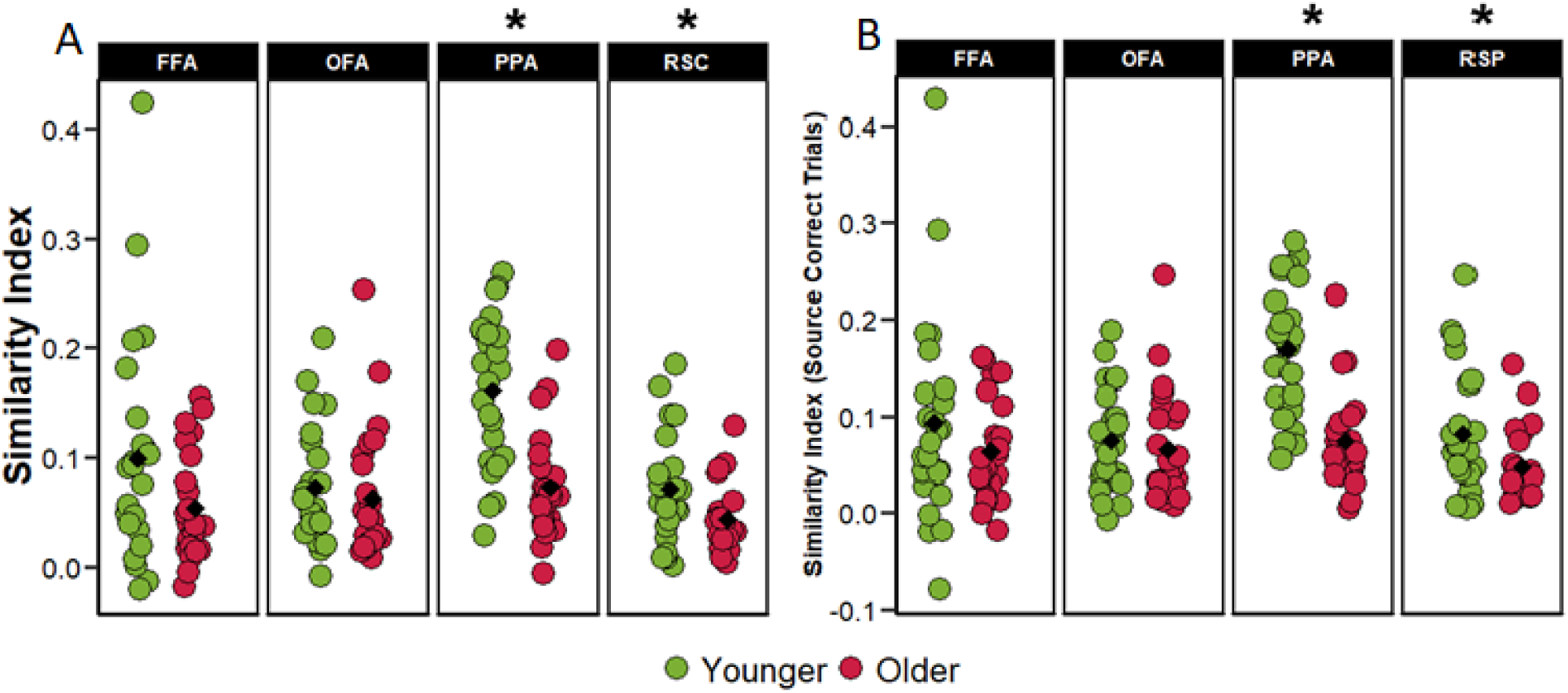
(A) Within – Between similarity indices computed collapsing across memory performance. (B) Within – Between similarity indices computed for only those trials that went on to receive a source-correct response at subsequent retrieval.

As with the analyses of the differentiation index, the within – between similarity indices were also computed separately for trials binned into two categories depending on whether or not the trial received a source-correct memory response at retrieval. A 2 (age group) x 4 (ROI) x 2 (memory status) mixed factorial ANOVA resulted in a main effect of age group (F_(1, 46)_ = 12.894, p < .001, partial-*η*^2^ = .219), a main effect of ROI (F_(2.34, 107.47)_ = 10.873, p < .001, partial-*η*^2^ = .191), an age-by-ROI interaction (F_(2.34, 107.47)_ = 4.480, p = .010, partial-*η*^2^ = .089), and a three-way interaction between age, ROI and memory status (F_(2.39, 109.99)_ = 3.542, p = .025, partial-*η*^2^ = .071). The analysis did not identify a main effect of memory (F_(1, 46)_ = 3.074, p = .098, partial-*η*^2^ = .063), nor any two-way interactions between memory and Age group or ROI (ps > 0.20). Subsequent pairwise comparisons demonstrated that the within-between similarity indices computed separately for the two classes of memory judgment were not significantly different from each other in either ROI in either age group (ps > 0.142).

For the reasons described above (Methods 2.5.6), we repeated the foregoing analyses using only those trials that went on to give rise to a correct source memory judgment, allowing an assessment of whether age-differences in pattern similarity were driven by age-differences in the number of successful memory trials contributing to the similarity metrics. A 2 (age group) x 4 (ROI) mixed ANOVA revealed significant main effects of age (F_(1, 46)_ = 12.071, p = 0.001, partial-*η*^2^ = 0.208), and ROI (F_(2.34, 107.43)_ = 10.550, p < 0.001, partial-*η*^2^ = 0.187), along with significant age by ROI interaction (F_(2.34, 107.43)_ = 5.325, p = 0.004, partial-*η*^2^ = 0.104). A follow-up ANOVA on the data for the scene-selective ROIs revealed significant main effects of age group (F_(1, 46)_ = 20.830, p < 0.001, partial-*η*^2^ = 0.312), and ROI (F_(1, 46)_ = 58.860, p < 0.001, partial-*η*^2^ = 0.561), as well as an age-by-ROI interaction (F_(1, 46)_ = 16.221, p < 0.001, partial-*η*^2^ = 0.261). ANOVA of the face-selective ROIs failed to identify any significant effects: age (F_(1,46)_ = 1.647, p = 0.206, partial-*η*^2^ = 0.035); ROI (F_(1,46)_ = 0.320, p = 0.574, partial-*η*^2^ = 0.007); age x ROI interaction (F_(1, 46)_ = 0.558, p = 0.459, partial-*η*^2^ = 0.012). As Figure 5.B illustrates, the similarity indices demonstrated age-related reductions in both the PPA and RSC (t_(41.62)_ = 5.543, p < 0.001, d = 1.600, and t_(37.12)_ = 2.328, p = 0.025, d = 0.672, respectively), while age effects were absent in the two face-selective ROIs (t_(33.53)_ = 1.23, p = 0.226, d = 0.356 and t_(45.54)_ = 0.575, p = 0.568, d = 0.166 in the FFA and OFA respectively). Similarity indices were however significantly different from zero in all ROIs and age groups (ps < 0.001). Thus, as with the differentiation index, when pattern similarity analysis was restricted to encoding trials associated with a correct subsequent source memory judgment robust age effects were evident in scene- but not face-selective ROIs.

### 3.4. Relationship between neural differentiation and subsequent memory performance

In light of prior findings (Koen et al., 2019), and as described in the methods, we ran a series of multiple regression analyses in which age group and the differentiation indices from each ROI were employed as predictors of subsequent source and item memory performance. As described in Methods, the initial multiple regression models included the ROI x Age interaction terms, however, in no case was the interaction significant (p > 0.05). Therefore, Table 6 presents the partial correlations between neural differentiation and performance after controlling for age group. As is evident from the table, the partial correlations between differentiation indices and source memory performance achieved significance only in the PPA (as would be predicted from the findings of Koen et al., 2019). This was the true both when computing the differentiation index collapsing across memory performance and when selecting only the source-correct trials. Moreover, these relationships between differentiation in the PPA and source memory performance remained significant after controlling for both age and item memory performance (collapsed across all trials: r_partial_ = 0.334, p = 0.023; source-correct trials: r_partial_ = 0.314, p = 0.033). The partial relationships controlling for age group are illustrated in Figure 6. Analogous analyses were conducted for the pattern similarity indices: no significant relationships between similarity indices and memory performance were identified (p > 0.092).

**Figure 6:**
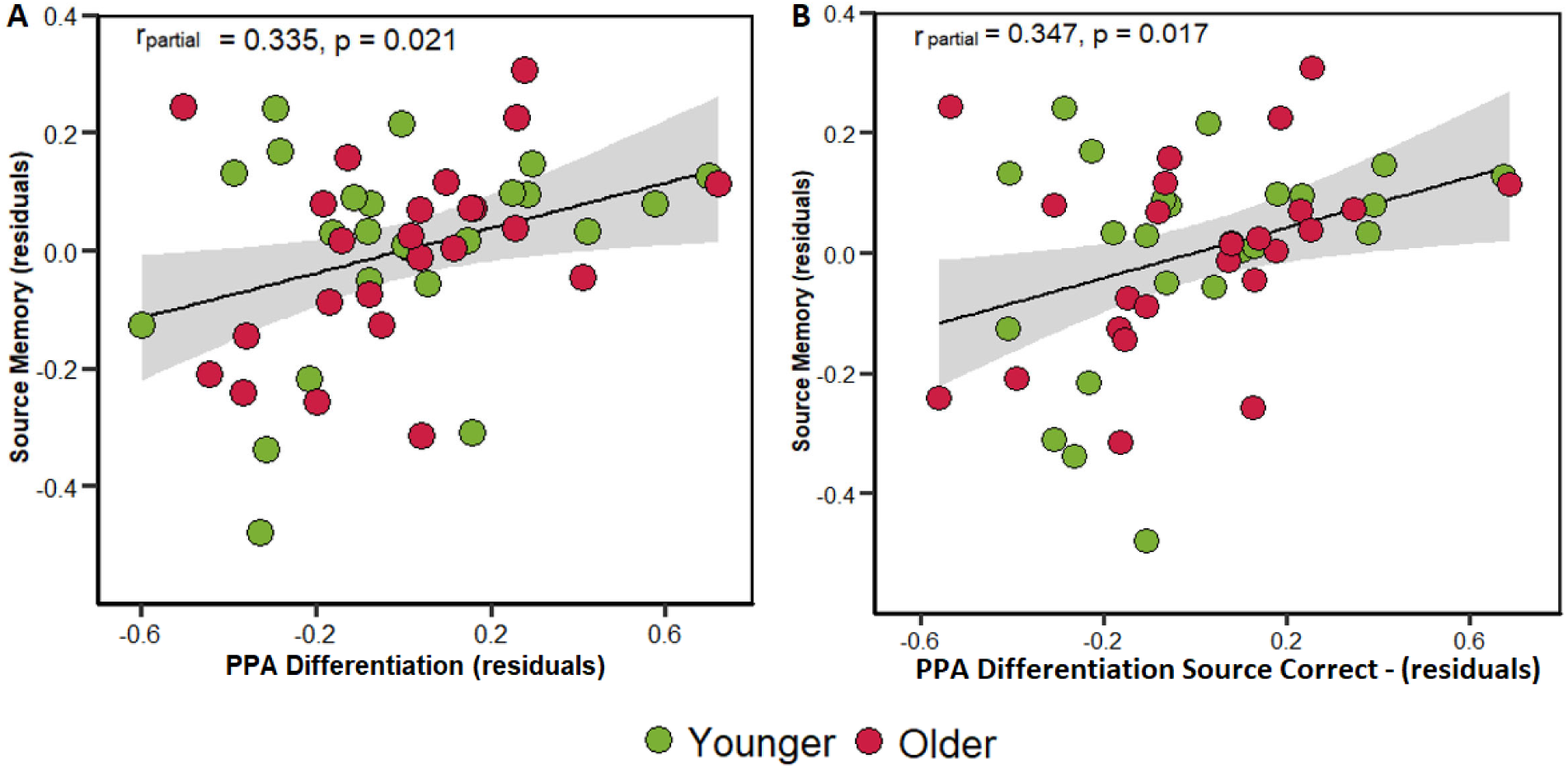
Scatterplots illustrating the partial correlations (controlling for age group) between PPA differentiation indices with source memory performance. Plot A illustrates the relationship between source memory and differentiation index collapsed across all encoding trials. Plot B illustrates the same relationship but restricted only to the trials that went on to receive a source correct memory response.

**Table 6:** Partial correlations (p-values) between item memory and source memory performance and differentiation index when controlling for age group. The differentiation indices were computed either across all encoding trials (first two columns) or only for those encoding trials that were associated with a source-correct memory response (second two columns).

|  | Collapsed across all trials |  | Source-correct trials |  |
| --- | --- | --- | --- | --- |
|  | Item Memory | Source Memory | Item Memory | Source Memory |
| *FFA* | -0.14 (0.33) | -0.12 (0.43) | -0.08 (0.58) | -0.01 (0.94) |
| *OFA* | 0.07 (0.64) | 0.08 (0.57) | 0.15 (0.32) | 0.08 (0.57) |
| *PPA* | 0.14 (0.34) | 0.34 (0.02) | 0.18 (0.22) | 0.35 (0.02) |
| *RSC* | 0.10 (0.52) | 0.15 (0.30) | 0.04 (0.80) | 0.10 (0.50) |

### 3.5. Relationship between neural differentiation and neuropsychological test performance

Given prior findings of a positive, age-invariant, relationship between the PPA differentiation index and the fluency component derived from the neuropsychological test battery (see Introduction), we examined whether a similar relationship was evident in the present study. When collapsed across all trials regardless of subsequent memory, the partial correlation (controlling for age) between the differentiation index and fluency factor scores was not significant in the PPA, r_partial_ = −0.009, p = 0.951 (nor in the RSC, r_partial_ = 0.112, p = 0.454). The relationship was also absent when the differentiation index was derived from source correct trials only, PPA: r_partial_ = 0.105, p = 0.482, RSC: r_partial_ = 0.170, p = 0.255.

Additional exploratory analyses were performed to examine the relationship between the other three cognitive constructs derived from the neuropsychological test battery (see Table 1) and the 4 ROIs. These analyses revealed an age-invariant relationship (controlling for age group) between the FFA differentiation index and the memory factor, r_partial_ = −0.430, p = 0.003, (p = 0.036 after Holmes-Bonferroni correction for multiple comparisons). The negative relationship between the FFA differentiation index and the memory factor remained significant when restricting the analysis to source correct trials, r_partial_ = −0.478, p < 0.001 (p = 0.003 after correction). We did not identify any significant relationships between pattern similarity metrics and neuropsychological test performance (p > 0.05).

## 4. Discussion

The current study employed a combination of univariate and multi-voxel analyses to examine age effects on category-level neural selectivity (or neural differentiation) during the encoding of images of faces and scenes prior to a subsequent memory test. Neural selectivity was examined in two scene- and two face-selective ROIs. The univariate and pattern similarity measures yielded convergent results indicating that scene-, but not face-selective, regions demonstrated reduced category-level selectivity with older age – that is, age-related neural dedifferentiation. The findings add to the already large literature describing age-related neural dedifferentiation effects (for review, see Koen and Rugg, 2019; Koen et al., 2019, 2020) and, importantly, also add to evidence suggesting that while the phenomenon is highly robust for scene stimuli, it is more elusive for other stimulus classes: faces in the present case, and objects in Koen et al. (2019). Additionally, analogous to the findings of Koen et al. (2019), the univariate metric of neural differentiation for scenes in the PPA demonstrated a positive, age-invariant, relationship with source memory performance.

Turning first to the behavioral findings, we observed no age differences either in study RT or in the vividness ratings assigned to the study items. Therefore, the age differences we identified in neural differentiation are unlikely to reflect the confounding effects of either of these variables. At test, younger adults outperformed their older counterparts in respect of both item and source memory performance, findings consistent with an extensive prior literature (for reviews, see Old & Naveh-Benjamin, 2008; Koen & Yonelinas, 2014). Given these age differences in memory performance, we examined neural differentiation indices derived not only from all experimental items (as in prior studies) but also from only those study trials attracting correct source judgments. The results of the two analyses revealed that the findings of age-related neural dedifferentiation in the scene-selective ROIs were not confounded by differential neural activity associated with successful vs. unsuccessful memory encoding.

Age-related reductions in neural specificity have been linked to cognitive declines associated with healthy aging (Koen & Rugg, 2019). This putative link is motivated by the notion that age-related weakening of dopaminergic neuromodulation results in reduced neural signal-to-noise and hence reduced specificity of neural representations (Li et al., 2001; Li & Rieckmann, 2014; see also Abdulrahman et al., 2017). The proposal that age-related neural dedifferentiation plays a role in cognitive decline receives further support from findings that dedifferentiation is associated with lower memory performance (Yassa et al., 2011; Berron et al., 2018; Bowman et al., 2019; Koen et al., 2019) and lower fluid processing ability (Park et al., 2010; Koen et al., 2019). However, although increasing age is undoubtedly associated with reduced neural selectivity, the existing evidence suggests that the relationship between neural differentiation and cognitive performance is not moderated by age, that is, it is age-invariant (Koen & Rugg, 2019). The present findings of an age-invariant relationship between scene differentiation in the PPA and subsequent source memory performance add to this evidence. These findings serve as a conceptual replication of those reported by Koen et al., (2019), although in that experiment, PPA differentiation was related more strongly to item than to source memory performance. This disparity likely reflects the different experimental procedures: whereas the category exemplars in the present study served as the contextual features targeted in the source memory test, in Koen et al. (2019) the exemplars were the test items themselves.

As noted in the Introduction, evidence for age-related neural dedifferentiation in the visual domain appears to be most consistent for scenes and faces. Thus, the present findings for scenes in the PPA and RSC are fully consistent with prior findings, whereas the null effects we report for faces in FFA and OFA run counter to several prior results (Park et al., 2004; Voss et al., 2008; Park et al., 2012; but see Payer et al. 2006). There are several factors that, either jointly or in combination, might account for these disparate findings. One factor concerns the presentation format of the stimuli. Whereas the faces in the present study were rendered in color, as best we can determine, prior studies reporting age-related differentiation for faces all employed gray-scale images. A second factor concerns the processing demands placed on the participants: whereas most prior studies reporting age effects on face specificity employed passive viewing conditions (Park et al., 2004; Voss et al., 2008; Park et al., 2010; but see Goh et al., 2010, and Burianová et al., 2013), here we employed a task that required active engagement with the experimental stimuli (as did Payer et al., 2006). If older adults have a greater tendency to “zone out” during passive viewing, the resulting reduction in attention to the experimental stimuli may manifest as reduced neural selectivity (see Koen et al., 2019, for a similar account of inconsistent findings for objects). Additionally, whereas prior studies reporting age-related differentiation typically employed blocked experimental designs, here we employed an event-related design in which different category exemplars were presented in an unpredictable order.

While some combination of the above-mentioned factors might account for the absence of age-related neural dedifferentiation for faces in the present study, they offer no insight into why dedifferentiation effects for scenes are so robust. Relevant to this question, a recent “lifetime experience hypothesis” (Koen & Rugg, 2019) posits that neural differentiation might be moderated by prior experience that accrues over the lifespan. The hypothesis proposes that accumulating lifetime experience facilitates the assimilation of novel category exemplars into perceptual schemas (Gilboa and Marlatte, 2017). If scene processing becomes increasingly schema-dependent with age, age-related neural dedifferentiation in scene ROIs might reflect more efficient assimilation of scene information into relevant schema(s). As was noted by Koen et al. (2019), this proposal receives support from their finding that age-related neural dedifferentiation in the PPA took the form of an age-related reduction in neural responses to scenes (neural attenuation), as here. By contrast, schemas for some other stimulus categories, such as canonical objects, high frequency words, and, possibly, faces, develop more rapidly and are largely fully formed by adolescence or early adulthood (Germine et al., 2011). By this view, therefore, the present findings of null age effects for face differentiation reflect the fact that young and older adults possess equally well-formed face schemas.

The mixed evidence for age differences in neural selectivity for different perceptual categories might also be explained by age differences in the perceptual processing of complex visual stimuli. For instance, age differences in neural differentiation may be more pronounced when viewing stimuli that comprise combinations of multiple, unpredictable features, such as scenes rather than faces. Notably, it has been reported that PPA activity is strongly modulated by scene complexity (Chai et al., 2010), whereby increasing complexity is associated with greater activity in the region (see Aminoff et al., 2013, for review). If, as has been suggested (e.g. Boutet et al., 2019; Meng et al., 2019), older adults are less able to differentiate visual detail, then age differences in neural selectivity in the PPA might be anticipated. In contrast, the null effects of age in neural selectivity for exemplars of canonical objects, words, or human faces, might reflect their relatively low visual complexity, along with, perhaps, higher schema congruency (see above).

For reasons that are presently unclear, we failed to replicate the finding (Koen et al., 2019) of a relationship between PPA differentiation and scores on a psychometric fluency factor. We did, however, identify a robust age-invariant negative relationship between the FFA differentiation index and the memory factor derived from the neuropsychological test scores. This finding should be treated with caution since it was unpredicted and, to our knowledge, is without precedent. On its face, however, it suggests that relatively greater neural selectivity does not invariably confer a cognitive benefit.

We note a number of limitations of the present study. First, measuring neural selectivity at the category level might not provide a sensitive enough measure to detect age differences in the fidelity of face (or object) representations, and it is possible that item-level measures would yield different findings (cf. Goh et al., 2010; St Laurent et al., 2014; Sommer et al., 2019; Trelle et al., 2019). Second, like prior studies of neural dedifferentiation, the present study employed a cross-sectional design. Hence, the reported age differences cannot unambiguously be attributed to the effects of aging (e.g. Rugg, 2016). Finally, it is unclear to what extent the present (and previous) findings reflect age differences the variability or the shape – as opposed to the gain (see Methods) - of stimulus-elicited hemodynamic responses (D’Esposito et al., 2003).

In conclusion, although increasing age is associated with reduced neural differentiation between different visual categories, the present study adds to the evidence that this is easier to demonstrate for visual scenes than for other visual categories. The findings also add to the evidence that neural differentiation can predict individual differences in cognitive performance throughout the lifespan. Whereas the functional significance of age-related neural dedifferentiation largely remains unclear, individual differences in neural differentiation appear to be determined by both age-dependent and age-invariant factors.

## References

Abdi H, Williams LJ (2010) *Principal Component Analysis*. In: Encyclopedia of ecology, Vol 2 (Jørgensen SE, Fath BD, eds), pp 2940–2949. Oxford: Elsevier.

Abdulrahman H, Fletcher PC, Bullmore E, Morcom AM (2017) Dopamine and memory dedifferentiation in aging. NeuroImage, 153, 211–220. https://doi.org/10.1016/j.neuroimage.2015.03.031

Aminoff EM, Kveraga K, Bar M (2013) The role of the parahippocampal cortex in cognition. Trends in Cognitive Sciences, 17(8), 379–390. https://doi.org/10.1016/j.tics.2013.06.009

Benton AL (1968) Differential behavioral effects in frontal lobe disease. Neuropsychologia 6:53– 60. https://doi.org/10.1016/0028-3932(68)90038-9

Berron D, Neumann K, Maass A, Schütze H, Fliessbach K, Kiven V, Jessen F, Sauvage M, Kumaran D, Düzel E (2018) Age-related functional changes in domain-specific medial temporal lobe pathways. Neurobiology of Aging, 65, 86–97. https://doi.org/10.1016/j.neurobiolaging.2017.12.030

Boutet I, Dawod K, Chiasson F, Brown O, Collin C (2019) Perceptual Similarity Can Drive Age-Related Elevation of False Recognition. Frontiers in Psychology, 10. https://doi.org/10.3389/fpsyg.2019.00743

Bowman CR, Chamberlain JD, Dennis NA (2019) Sensory Representations Supporting Memory Specificity: Age Effects on Behavioral and Neural Discriminability. The Journal of Neuroscience: The Official Journal of the Society for Neuroscience, 39(12), 2265–2275. https://doi.org/10.1523/JNEUROSCI.2022-18.2019

Burianová H, Lee Y, Grady CL, Moscovitch M (2013) Age-related dedifferentiation and compensatory changes in the functional network underlying face processing. Neurobiology of Aging, 34(12), 2759–2767. https://doi.org/10.1016/j.neurobiolaging.2013.06.016

Carp J, Park J, Polk TA, Park DC (2011) Age differences in neural distinctiveness revealed by multi-voxel pattern analysis. NeuroImage, 56(2), 736–743. https://doi.org/10.1016/j.neuroimage.2010.04.267

Chai XJ, Ofen N, Jacobs LF, Gabrieli JDE (2010). Scene complexity: Influence on perception, memory, and development in the medial temporal lobe. Frontiers in Human Neuroscience, 4. https://doi.org/10.3389/fnhum.2010.00021

Chee MWL, Goh JOS, Venkatraman V, Tan JC, Gutchess A, Sutton B, Hebrank A, Leshikar E, Park D (2006). Age-related Changes in Object Processing and Contextual Binding Revealed Using fMR Adaptation. Journal of Cognitive Neuroscience, 18(4), 495–507. https://doi.org/10.1162/jocn.2006.18.4.495

Cohen J (1988) Statistical power analysis for the social sciences, Ed 2. Hillsdale, NJ: Erlbaum.

de Chastelaine M, Wang TH, Minton B, Muftuler LT, Rugg MD (2011) The effects of age, memory performance, and callosal integrity on the neural correlates of successful associative encoding. Cerebral Cortex (New York, N.Y.: 1991), 21(9), 2166–2176. https://doi.org/10.1093/cercor/bhq294

de Chastelaine M, Mattson JT, Wang TH, Donley BE, Rugg MD (2015) Sensitivity of negative subsequent memory and task-negative effects to age and associative memory performance. Brain Research, 1612, 16–29. https://doi.org/10.1016/j.brainres.2014.09.045

de Chastelaine M, Mattson JT, Wang TH, Donley BE, Rugg MD (2016) The relationships between age, associative memory performance, and the neural correlates of successful associative memory encoding. Neurobiology of Aging, 42, 163–176. https://doi.org/10.1016/j.neurobiolaging.2016.03.015

Delis DC, Kramer JH, Kaplan E, Ober BA (2000) California verbal learning test, Ed 2. San Antonio, TX: The Psychological Corporation.

D’Esposito M, Deouell LY, Gazzaley A (2003) Alterations in the BOLD fMRI signal with ageing and disease: A challenge for neuroimaging. Nature Reviews Neuroscience, 4(11), 863–872. https://doi.org/10.1038/nrn1246

Du Y, Buchsbaum BR, Grady CL, Alain C (2016) Increased activity in frontal motor cortex compensates impaired speech perception in older adults. Nature Communications, 7(1), 12241. https://doi.org/10.1038/ncomms12241

Germine LT, Duchaine B, Nakayama K (2011) Where cognitive development and aging meet: Face learning ability peaks after age 30. Cognition, 118(2), 201–210. https://doi.org/10.1016/j.cognition.2010.11.002

Gilboa A, Marlatte H (2017) Neurobiology of Schemas and Schema-Mediated Memory. Trends in Cognitive Sciences, 21(8), 618–631. https://doi.org/10.1016/j.tics.2017.04.013

Goh JO, Suzuki A, Park DC (2010) Reduced neural selectivity increases fMRI adaptation with age during face discrimination. NeuroImage, 51(1), 336–344. https://doi.org/10.1016/j.neuroimage.2010.01.107

Gottlieb LJ, Uncapher MR, Rugg MD (2010) Dissociation of the neural correlates of visual and auditory contextual encoding. Neuropsychologia, 48(1), 137–144. https://doi.org/10.1016/j.neuropsychologia.2009.08.019

Greenhouse SW, Geisser S (1959) On methods in the analysis of profile data. Psychometrika 24:95–112. https://doi.org/10.1007/BF02289823

Hotelling H (1933) Analysis of a complex of statistical variables into principal components. Journal of Educational Psychology, 24(6), 417–441. https://doi.org/10.1037/h0071325

Kim S (2015) ppcor: An R Package for a Fast Calculation to Semi-partial Correlation Coefficients. Communications for Statistical Applications and Methods, 22(6), 665–674. https://doi.org/10.5351/CSAM.2015.22.6.665

Koen JD, Hauck N, Rugg MD (2019) The Relationship between Age, Neural Differentiation, and Memory Performance. The Journal of Neuroscience: The Official Journal of the Society for Neuroscience, 39(1), 149–162. https://doi.org/10.1523/JNEUROSCI.1498-18.2018

Koen JD, Horne ED, Hauck N, Rugg, MD (2018) Age-related Differences in Prestimulus Subsequent Memory Effects Assessed with Event-related Potentials. Journal of Cognitive Neuroscience, 30(6), 829–850. https://doi.org/10.1162/jocn_a_01249

Koen JD, Rugg MD (2019) Neural Dedifferentiation in the Aging Brain. Trends in Cognitive Sciences, 23(7), 547–559. https://doi.org/10.1016/j.tics.2019.04.012

Koen JD, Yonelinas AP (2014) The effects of healthy aging, amnestic mild cognitive impairment, and Alzheimer’s disease on recollection and familiarity: A meta-analytic review. Neuropsychology Review, 24(3), 332–354. https://doi.org/10.1007/s11065-014-9266-5

Koen JD, Srokova S, Rugg MD (2020) Age-Related Neural Dedifferentiation and Cognition, *Current Opinion in Behavioral Sciences*. (in press)

Kriegeskorte N, Mur M, Bandettini PA (2008) Representational similarity analysis—Connecting the branches of systems neuroscience. Frontiers in Systems Neuroscience, 2. https://doi.org/10.3389/neuro.06.004.2008

Li S-C, Lindenberger U, Sikström S (2001) Aging cognition: From neuromodulation to representation. Trends in Cognitive Sciences, 5(11), 479–486. https://doi.org/10.1016/S1364-6613(00)01769-1

Li S-C, Rieckmann A (2014) Neuromodulation and aging: Implications of aging neuronal gain control on cognition. Current Opinion in Neurobiology, 29, 148–158. https://doi.org/10.1016/j.conb.2014.07.009

Mattson JT, Wang TH, de Chastelaine M, Rugg MD (2014) Effects of Age on Negative Subsequent Memory Effects Associated with the Encoding of Item and Item–Context Information. Cerebral Cortex (New York, NY), 24(12), 3322–3333. https://doi.org/10.1093/cercor/bht193

Meng Q, Wang B, Cui D, Liu N, Huang Y, Chen L, Ma Y (2019) Age-related changes in local and global visual perception. Journal of Vision, 19(1), 10. https://doi.org/10.1167/19.1.10

Minear M, Park DC (2004) A lifespan database of adult facial stimuli. Behavior Research Methods, Instruments, & Computers, 36(4), 630–633. https://doi.org/10.3758/BF03206543

Mumford JA, Davis T, Poldrack RA (2014) The impact of study design on pattern estimation for single-trial multivariate pattern analysis. NeuroImage, 103, 130–138. https://doi.org/10.1016/j.neuroimage.2014.09.026

Old SR, Naveh-Benjamin M (2008) Differential effects of age on item and associative measures of memory: A meta-analysis. Psychology and Aging, 23(1), 104–118. https://doi.org/10.1037/0882-7974.23.1.104

Park DC, Polk TA, Park R, Minear M, Savage A, Smith MR (2004) Aging reduces neural specialization in ventral visual cortex. Proceedings of the National Academy of Sciences, 101(35), 13091–13095. https://doi.org/10.1073/pnas.0405148101

Park J, Carp J, Hebrank A, Park DC, Polk TA (2010) Neural specificity predicts fluid processing ability in older adults. The Journal of Neuroscience: The Official Journal of the Society for Neuroscience, 30(27), 9253–9259. https://doi.org/10.1523/JNEUROSCI.0853-10.2010

Park J, Carp J, Kennedy KM, Rodrigue KM, Bischof GN, Huang C-M, Rieck JR, Polk TA, Park DC (2012) Neural broadening or neural attenuation? Investigating age-related dedifferentiation in the face network in a large lifespan sample. The Journal of Neuroscience: The Official Journal of the Society for Neuroscience, 32(6), 2154–2158. https://doi.org/10.1523/JNEUROSCI.4494-11.2012

Payer D, Marshuetz C, Sutton B, Hebrank A, Welsh RC, Park DC (2006) Decreased neural specialization in old adults on a working memory task: NeuroReport, 17(5), 487–491. https://doi.org/10.1097/01.wnr.0000209005.40481.31

R Core Team (2017) R: a language and environment for statistical computing. Vienna, Austria: R Foundation.

Raven J, Raven JC, Courth JH (2000) Manual for Raven’s progressive matrices and vocabulary scales. Section 4: the advanced progressive matrices. San Antonio, TX: Harcourt Assessment.

Reitan RM, Wolfson D (1985) The Halstead-Reitan neuropsychological test battery: therapy and clinical interpretation. Tucson, AZ: Neuropsychological.

Revelle WR (2017) psych: procedures for psychological, psychometric, and personality research. Vienna, Austria: R Foundation.

Rissman J, Gazzaley A, D’Esposito M (2004) Measuring functional connectivity during distinct stages of a cognitive task. NeuroImage, 23(2), 752–763. https://doi.org/10.1016/j.neuroimage.2004.06.035

Rugg MD (2016) Interpreting Age-Related Differences in Memory-Related Neural Activity. Oxford University Press. https://www.oxfordscholarship.com/view/10.1093/acprof:oso/9780199372935.001.0001/acprof-9780199372935-chapter-8

Singmann H, Bolker B, Westfall J, Aust F (2016) afex: analysis of factorial experiments. Vienna, Austria: R Foundation.

Smith A (1982) Symbol digit modalities test (SDMT) manual. Los Angeles: Western Psychological Services.

Snodgrass JG, Corwin J (1988). Pragmatics of measuring recognition memory: applications to dementia and amnesia. J Exp Psychol Gen 117:34 –50. https://doi.org/10.1037/0096-3445.117.1.34

Spreen O, Benton AL (1977) Neurosensory center comprehensive examination for aphasia. Victoria, BC, Canada: Neuropsychology Laboratory.

Sommer VR, Fandakova Y, Grandy TH, Shing YL, Werkle-Bergner M, Sander MC (2019) Neural Pattern Similarity Differentially Relates to Memory Performance in Younger and Older Adults. Journal of Neuroscience, 39(41), 8089–8099. https://doi.org/10.1523/JNEUROSCI.0197-19.2019

St-Laurent M, Abdi H, Bondad A, Buchsbaum BR (2014) Memory Reactivation in Healthy Aging: Evidence of Stimulus-Specific Dedifferentiation. Journal of Neuroscience, 34(12), 4175–4186. https://doi.org/10.1523/JNEUROSCI.3054-13.2014

Thakral PP, Wang TH, Rugg MD (2019) Effects of age on across-participant variability of cortical reinstatement effects. NeuroImage, 191, 162–175. https://doi.org/10.1016/j.neuroimage.2019.02.005

Torchiano M (2019) effsize: Efficient Effect Size Computation. https://zenodo.org/record/1480624

Trelle AN, Henson RN, Simons JS (2019) Neural evidence for age-related differences in representational quality and strategic retrieval processes. Neurobiology of Aging. https://doi.org/10.1016/j.neurobiolaging.2019.07.012

Voss MW, Erickson KI, Chaddock L, Prakash RS, Colcombe SJ, Morris KS, Doerksen S, Hu L, McAuley E, Kramer AF (2008) Dedifferentiation in the visual cortex: An fMRI investigation of individual differences in older adults. Brain Research, 1244, 121–131. https://doi.org/10.1016/j.brainres.2008.09.051

Wang TH, Johnson JD, de Chastelaine M, Donley BE, Rugg MD (2016) The Effects of Age on the Neural Correlates of Recollection Success, Recollection-Related Cortical Reinstatement, and Post-Retrieval Monitoring. Cerebral Cortex (New York, N.Y.: 1991), 26(4), 1698–1714. https://doi.org/10.1093/cercor/bhu333

Wechsler D (1981) WAIS-R: Wechsler adult intelligence scale-revised. New York: The Psychological Corporation.

Wechsler D (2001) Wechsler test of adult reading. San Antonio, TX: The Psychological Corporation.

Wechsler D (2009) Wechsler memory scale, 4th ed. San Antonio, TX: The Psychological Corporation.

Yarkoni T, Poldrack RA, Nichols TE, Van Essen DC, Wager TD (2011). Large-scale automated synthesis of human functional neuroimaging data. Nature Methods, 8(8), 665–670. https://doi.org/10.1038/nmeth.1635

Yassa MA, Mattfeld AT, Stark SM., Stark CEL (2011) Age-related memory deficits linked to circuit-specific disruptions in the hippocampus. Proceedings of the National Academy of Sciences of the United States of America, 108(21), 8873–8878. https://doi.org/10.1073/pnas.1101567108

Zheng L, Gao Z, Xiao X, Ye Z, Chen C, Xue G (2018) Reduced Fidelity of Neural Representation Underlies Episodic Memory Decline in Normal Aging. Cerebral Cortex, 28(7), 2283–2296. https://doi.org/10.1093/cercor/bhx130

